# FiPhA: An Open-Source Platform for Fiber Photometry Analysis

**DOI:** 10.1101/2023.07.21.550098

**Authors:** Matthew F. Bridge, Leslie R. Wilson, Sambit Panda, Korey D. Stevanovic, Ayland C. Letsinger, Sandra McBride, Jesse D. Cushman

## Abstract

**Significance:** Fiber photometry is a widely used technique in modern behavioral neuroscience, employing genetically encoded fluorescent sensors to monitor neural activity and neurotransmitter release in awake-behaving animals, However, analyzing photometry data can be both laborious and time-consuming.

**Aim:** We propose the FiPhA (Fiber Photometry Analysis) app, which is a general-purpose fiber photometry analysis application. The goal is to develop a pipeline suitable for a wide range of photometry approaches, including spectrally resolved, camera-based, and lock-in demodulation.

**Approach:** FiPhA was developed using the R Shiny framework and offers interactive visualization, quality control, and batch processing functionalities in a user-friendly interface.

**Results:** This application simplifies and streamlines the analysis process, thereby reducing labor and time requirements. It offers interactive visualizations, event-triggered average processing, powerful tools for filtering behavioral events and quality control features.

**Conclusions:** FiPhA is a valuable tool for behavioral neuroscientists working with discrete, event-based fiber photometry data. It addresses the challenges associated with analyzing and investigating such data, offering a robust and user-friendly solution without the complexity of having to hand-design custom analysis pipelines. This application thus helps standardize an approach to fiber photometry analysis.

## 1 Introduction

Monitoring neurotransmission in the brain of freely moving animals is important to understand how real-time neuronal activity and specific behavioral events are correlated. Identifying this relationship can help elucidate the underling circuitry of various brain regions, and how that circuitry is manipulated by various toxins, diseases, etc. One way to monitor neurotransmission is fiber photometry (FP), which is an optical technique used for recording cellular activity by detecting bulk fluorescence signals within cell populations or brain regions.^1^ Cells are fluorescently labeled using viral vectors or trans-genetic approaches either at cell bodies^2^ or within axon terminals.^3^ While FP was first developed as a tool to measure intracellular calcium,^4–6^ it has recently been used to detect the binding of specific neurotransmitters to fluorescently labeled post synaptic receptors.^7–10^ Recording within multiple brain regions simultaneously is also possible with fiber photometry due to the relatively small size of the implantable optical probes and lightweight fiber cables.^7, 11^

There has been increased interest in the use of fiber photometry for neuroscience, and researchers primarily depend on lock-in demodulation systems or other custom setups. These systems produce an immense amount of data, which were conventionally analyzed using custom MATLAB or Python scripts. Since then, several open-source measurement tools and analysis packages have been developed that operate on all major operating systems and commonly used programming languages.^12–14^ These packages have helped standardize fiber photometry analysis, but suffer from a few issues: 1) these packages are optimized for lock-in demodulation systems and have either limited or absent support for other systems, 2) they can easily analyze behavioral triggered transistor-transistor-logic (TTL) pulses, but require additional data preprocessing to analyze experiments that have non-behaviorally triggered events, and 3) they can have unintuitive and unappealing user-interfaces and visualizations.

We introduce the FiPhA app, which enables researchers to analyze behavioral data and fiber photometry data within the same user interface. FiPhA makes it possible for users to import multiple data formats from many photometry systems currently available on the market. We have validated that FiPhA can analyze data from a lock-in demodulation system,^15^ a camera-based system,^16^ and a spectrally resolved system,^17^ allowing users to flexibly define fluorescent activity during events and intervals of interest. FiPhA can also easily incorporate behavioral data collected using commercially available tracking systems. An important benefit of using FiPhA for analysis is that the data acquired needs little to no preprocessing before being imported into the app. Additionally, FiPhA can not only quickly align behavioral and fiber photometry data but also offers aesthetically pleasing visualizations. FiPhA thus helps standardize an approach to fiber photometry analysis that is both robust and user-friendly.

## 2 General Overview of FiPhA

FiPhA was developed using RStudio (v2022.x) and R (v4.2.x). It uses the Shiny framework to provide user interface functionality and the Plotly package for its interactive data visualizations. A pipeline for jointly analyzing behavioral and photometry data using the FiPhA app is illustrated in Fig. 1. This pipeline contains common steps for analyzing behavioral and photometry data in tandem, including preprocessing data, creating event-triggered averages, filtering behavioral events, and execution of various data analysis tasks. Analysis functionality includes the calculation of mean event-triggered averages, summary statistics of event intervals and their visualization in box-and-whisker plots, and graphics depicting of fluorescence activity by event across time in the form of heatmaps. The data can be saved as a “.rds” file for future analysis and the analysis of the event-triggered averages can also be exported as a “.xlsx” file.

**Figure 1:**
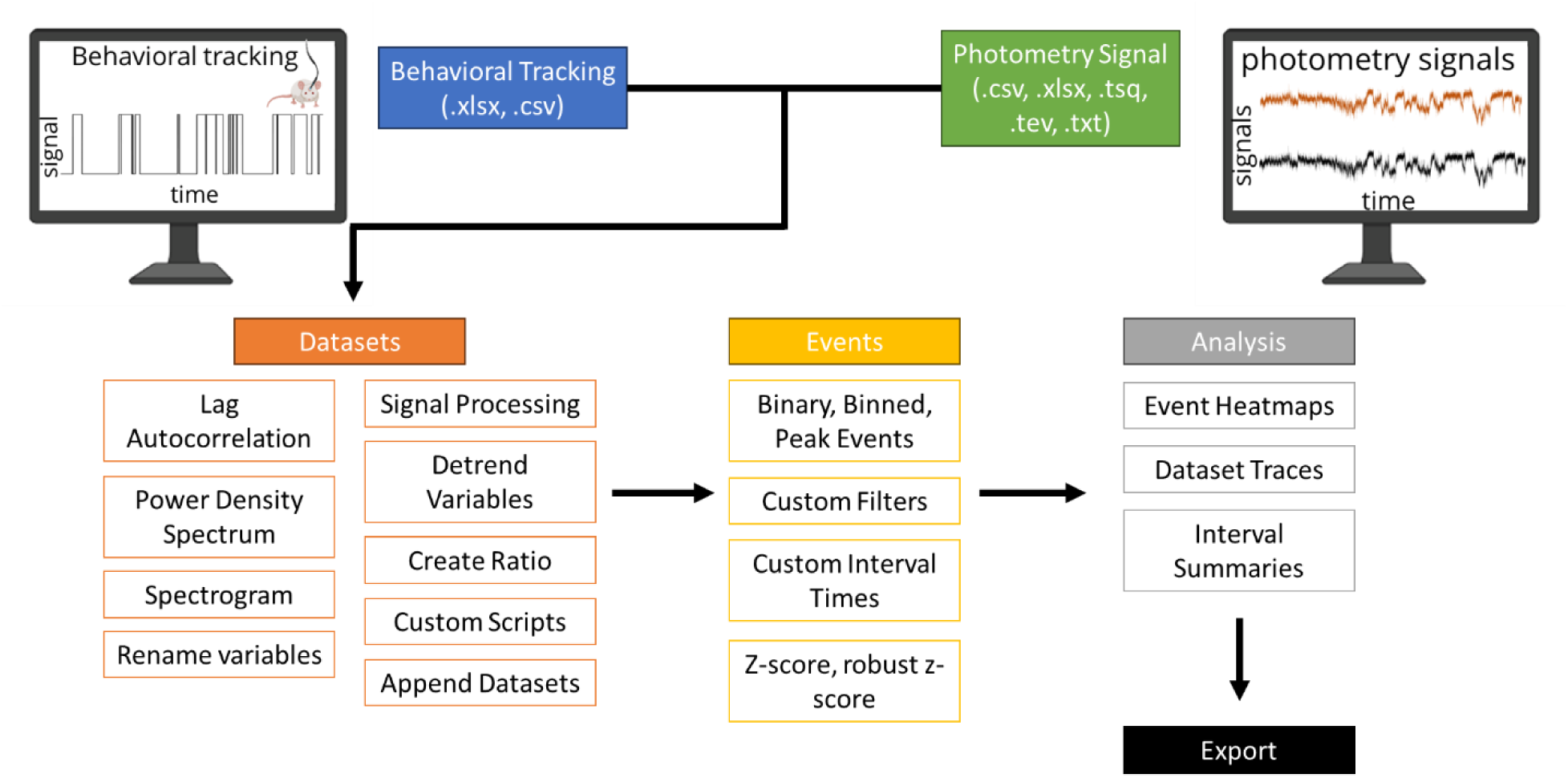
Typical workflow for analysis with the FiPhA app.

### 2.1 Datasets

#### 2.1.1 Importing Recordings

During fiber photometry experiments, experimenters typically collect photometry signal and behavioral tracking data using separate software packages. The FiPhA app allows importing data from various formats (.xlsx, .csv, .txt, .tsq, etc.). Prior to analysis, photometry signals collected using spectrally resolved photometry systems necessarily undergo a decomposition into signals of interest through the application of either a linear unmixing algorithm or a “summary statistic” based option with user-defined ranges of wavelengths for optimal signal identification when using fluorophores with overlapped emission spectra (Fig. 2). FiPhA simplifies the linear unmixing step by allowing users to import the raw spectrometer file, set the selected wavelength row, data row, and collection frequency, and then import a specific spectrometer reference file within a single interface. This spectrometer reference file contains standard fluorescence spectrograms specific to the wavelengths of the experimentally collected fluorescence signals.

**Figure 2:**
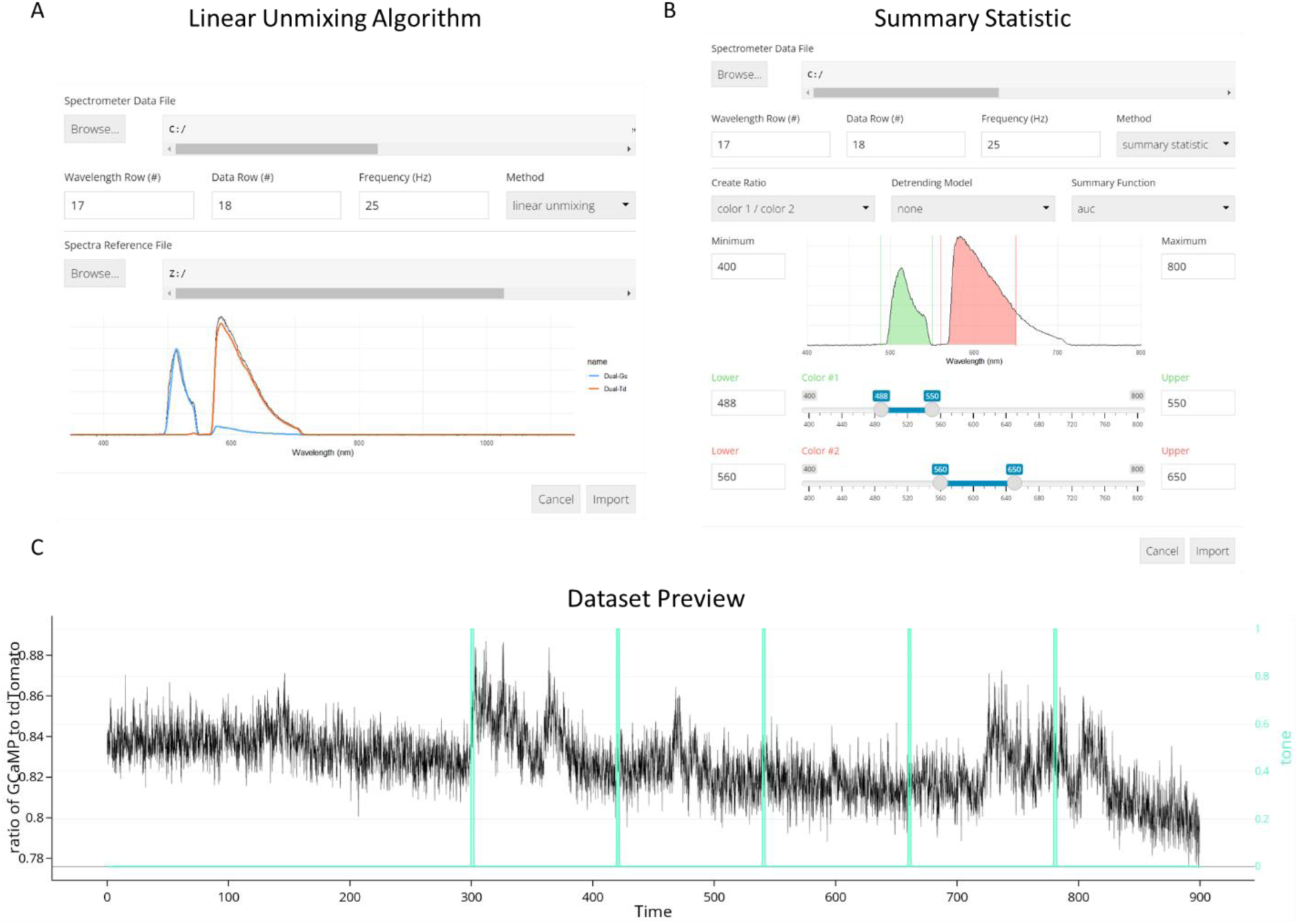
Import options for data collected using spectrally resolved photometry systems. (A) The linear unmixing algorithm showing a GCaMP and tdTomato fluorescence peak as well as a reference spectrum. (B) Importing data using the summary statistic option with calculating the area under the curve. (C) Dataset Preview of a 900 second photometry recording session aligned with TTL pulses of 5 tone presentations.

The “summary statistic” option allows for the creation of a ratio of two fluorescence signals derived from user-specified wavelength ranges via application of a summary function (i.e. area under the curve (AUC), mean, and median), followed by automatic signal detrending. Spectrometer and behavioral data can then be plotted in the Preview tab located within the Datasets view.

Alternative fiber photometry data that are not spectrally-resolved can be imported into the FiPhA app. Data collected from the lock-in demodulation systems are typically saved in “.tsq” or “.tev” formats, which can be imported into the FiPhA app using the Fiber Photometry Gizmo located under the import tab drop-down menu. Users specify the “.tsq” or “.tev” files by selection of a target directory and the desired fluorescence channel using the “available streams” drop down menu.

Some fiber photometry collection systems will save the raw data as a tabular dataset in a “.xlsx” or “.csv” format. These can be easily imported into the FiPhA app using the Tabular Dataset function under the import tab drop-down menu. To use this feature, the header row numbers, data row numbers, collection frequencies, and workbook sheet number will need to be specified. Data can then be similarly visualized using the Preview tab in the Datasets view.

Previous analysis sessions using the FiPhA app can be saved as a “.rds” file and reimported using the previous session import option in the drop-down menu.

#### 2.1.2 Preview

After the data are imported into the FiPhA app, the recordings can be observed in the Preview tab. Additional recordings can then be added by simply selecting another recording to import, which then appear as a list under the top drop-down menu. In the Preview tab, the recording time is typically depicted on the x-axis, but can be changed on the left drop-down menu. Manipulating the y-axes can be done under the right drop-down menu. Two datasets can be viewed at once using the left and right y-axes.

#### 2.1.3 Signal Analysis

Signal analysis and investigation of periodic components of a given recording’s continuous variables can be performed using the Signal Analysis tab. Selection of a dataset will produce a lag autocorrelation plot, a power density spectrum plot, and a spectrogram along with options to customize the resulting figures. Lag-N autocorrelation is a measure of a signal’s correlation with a delayed (by N time points) copy of itself, while the power density spectrum and spectrogram both visualize the overall individual frequencies that contribute to a signal and how these change over time.

#### 2.1.4 Transforms

Recordings can be combined with additional datasets from external hardware (such as behavior information derived from video recordings) after a dataset has been imported. This functionality is available under the Transform menu. Alignment of datasets which do not begin simultaneously or those which have been recorded at different collection frequencies are addressed by nearest neighbor sampling of the behavioral dataset and concatenation of the resulting data as a new column.

Additional operations available under the Transform menu include the ability to append two datasets (row-wise), renaming either the dataset itself or any of its constituent variables, and calculating a ratio of two dataset variables Low-pass filters can be applied in tandem with down sampling of larger datasets. Linear scaling transformations can also be applied, which helps adjust the magnitude of a variable to that of another and is a necessary step in recreating the “robust z-score” transformation (see Sec. 2.2.2).

Photobleaching/photoswitching is a common phenomenon seen in fiber photometry experiments whereby the magnitude of fluorescent activity decreases over time. To account for this effect, activity can be modeled as a decreasing function of time via linear or exponential models. Model parameters may be either manually specified or estimated using least squares fits, and model residuals and predicted values can be returned.

#### 2.1.5 Custom Scripts

Users familiar with the R programming language may optionally apply custom transformations to imported datasets before event detection. Snippets of code are executed in an environment that has been set up to contain the dataset as a “data.table” object, an extension of the standard “data.frame” class in base R. Any transformations made are then carried over to the imported dataset. This functionality allows for the inclusion of complex operations in workflows that are not already features of FiPhA.

### 2.2 Events

#### 2.2.1 Binary, Binned, Peak Events

##### Binary events

An event series may be defined using a binary (TTL-like) indicator variable contained within a dataset. Events of this type will correspond to continuous periods which begin at each rising edge transition (0 to 1) through to the following falling edge (1 to 0). An inverted type is also available which performs the same identification strategy but instead with the logical complement of the specified variable.

##### Binned events

In some experimental setups, it may be desirable to divide a time series into equally sized bins for tasks such as peak counting. This type is parameterized by a starting time ***t*** and bin length ***L***. Beginning at ***t***, each successive interval of length ***L*** corresponds to a new event.

##### Peak events

Events may be identified using a peak detection method that uses a moving window of length ***L*** to identify periods of a continuous variable that exceed a user-defined threshold in terms of the number of standard deviations away from the window’s mean value.

##### Timestamped events

For certain experimental setups, events may not be suitably described using variables within the same dataset. In such cases, FiPhA enables users to manually input a list of start/end timestamps which correspond to the desired events.

#### 2.2.2 Normalizations

The FiPhA app provides the choice of three normalization methods based on the distributional properties of the signal.

##### Z-Score

The z-score is calculated by subtracting the mean of the interval of interest from the observation and dividing by the standard deviation. This normalization is best suited to normally distributed, uncorrelated observations.

##### ΔF/F

This normalization is commonly used when processing fiber photometry data and is the percent change in intensity relative to a baseline period. Here, the standard deviation is replaced by the average value. Because the average may be sensitive to large values, the ΔF/F normalization may not be appropriate for noisy signals.

##### Robust Z-Score

A more robust z-score is calculated by subtracting the median percent change in intensity of the interval of interest from the observation and dividing by the median absolute deviation.^12^ This method is less sensitive to outlying observations than the standard z-score, but may be difficult to interpret since its reference period is no longer centered at zero.

#### 2.2.3 Custom Filters

##### Conditions

Potential events may be conditioned on other variables within a dataset by specifying a valid R expression (e.g. ‘(time)’ < 360 & cond == T) that returns a logical value. This expression is evaluated within the context of the dataset for each time step, controlling the state of inclusion for any potential events prior to filtering. Any valid R function may also be used in the statement, and an option is available to control whether a potential event must meet the specified conditions over its entire length, or more loosely at one or more points within its length.

##### Filters

An event series definition may also include additional filters, which successively modifies the list of potential events depending on the signal type. Spurious events introduced by noisy measurements (such as those derived from video recordings of animal behavior) may be removed using some combination of these filters:

- Exclusion/inclusion of the first or last events.
- Temporal shifting of events by a constant time.
- Padding of events such that they meet a minimum length.
- Restriction of events to a maximum length of time.
- Exclusion of events that do not meet a specified minimum length.
- Exclusion of events that exceed a maximum length.
- Aggregation of events which occur within a specified amount of time of one another.
- Coalescing of overlapping events.
- Inclusion based on a minimum rate of occurrence.
- Exclusion/inclusion of events based on occurrence before/after a given timestamp.
- Exclusion of successive events which occur within a given time of one another.

#### 2.2.4 Custom Interval Times

Intervals of interest can be manually specified in a table, where each entry specifies the name, reference point, and start/end times in relation to the reference point. Available choices include the beginning of the event signal, the end of the event signal, the raw event signal itself, or a “beginning” variant that ensures no overlap occurs with preceding events or their intervals by adjusting the interval’s start time if necessary.

### 2.3 Analysis

The Analysis tab within FiPhA enables users to create several different visualizations of processed event series. Each plot highlights various features of the resulting data, allowing users to investigate the full range of events in several different contexts.

#### 2.3.1 Event Heat Maps

Heatmaps which visualize all processed events are produced under the Analysis tab. Events are aligned such that their event signals all start at time zero, with options to sort events by their order, total length, or area under the curve between either the full event or two specified points.

#### 2.3.2 Dataset Traces

Likewise, visualizations of averaged event responses are also available using the Analysis tab. For a given dataset, multiple event series may be plotted simultaneously with options to also include a shaded area that represents the mean plus or minus a user-provided number of standard deviations (or standard errors).

#### 2.3.3 Interval summaries

Boxplots containing summary values of event interval across multiple dataset and event series can be found under the Analysis tab. In addition to the mean interval value, the median interval value can also be plotted, as well the area under the curve value between two time points of each event.

### 2.4 Export

The Export tab within FiPhA allows for production of Excel workbooks, which contain all processed event data. Individual tabs corresponding to each dataset and columns for each event are included. By default, these notebooks are more machine-readable to ensure compatibility with software that deals with post-processed event data; however, several options exist to control the formatting of the resulting workbooks and produce more human-readable files. Response variable names, interval labels, and numerous timestamps (raw dataset time, time within each interval, and time relative to the start of an event signal) can also be included. Alignment of the resulting columns can also be done across events by time or interval type to assist in the comparison of events within Excel. It is also possible to save the active session as an R object, which can later be re-imported or manipulated in R without needing to use FiPhA.

## 3 Example Workflow

The FiPhA application can analyze data without an associated behavioral trigger. Here, an experiment was performed where ventral hippocampal acetylcholine levels during wheel running in mice was analyzed before and after receiving an intraperitoneal dose of clozapine n-oxide to alter the activity of a physical activity-related neural circuit. Two separate recordings were taken (before and after the intraperitoneal dosing) and scored using EthoVision XT 16 (Fig. 7A). With FiPhA, these separate recordings can be combined and jointly analyzed (Fig. 7B). Event triggered averages can be computed, where individual events are filtered, and interval lengths specified (Fig. 7C). This experiment uses behaviorally triggered events, unlike data from previous figures which examined stimuli presented by the experimenter, so there are various lengths for each wheel running event. The ΔF/F normalization was then performed, and two different views of the event triggered averages were visualized: 1) a trace of the normalized average and deviation across all events (Fig. 7D) and 2) a chronological heatmap of each normalized wheel running event over time (Fig. 7E). This provides an example of the flexibility with which FiPhA can be used for event triggered averaging for behaviorally triggered events.

**Figure 3:**
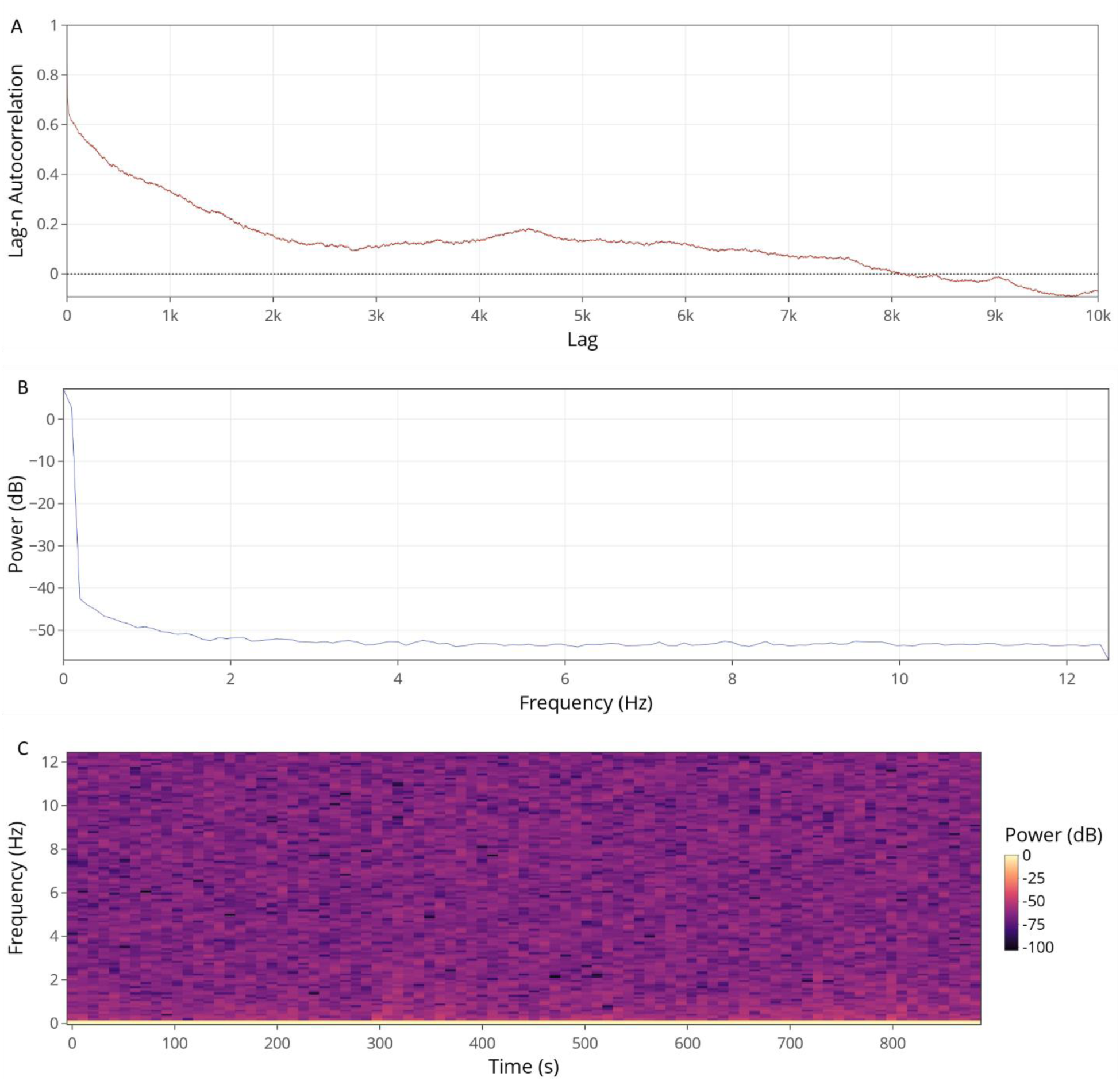
Functions located within the Signal Analysis tab (A), power density spectrum (B), lag autocorrelation (C), and spectrogram.

**Figure 4:**
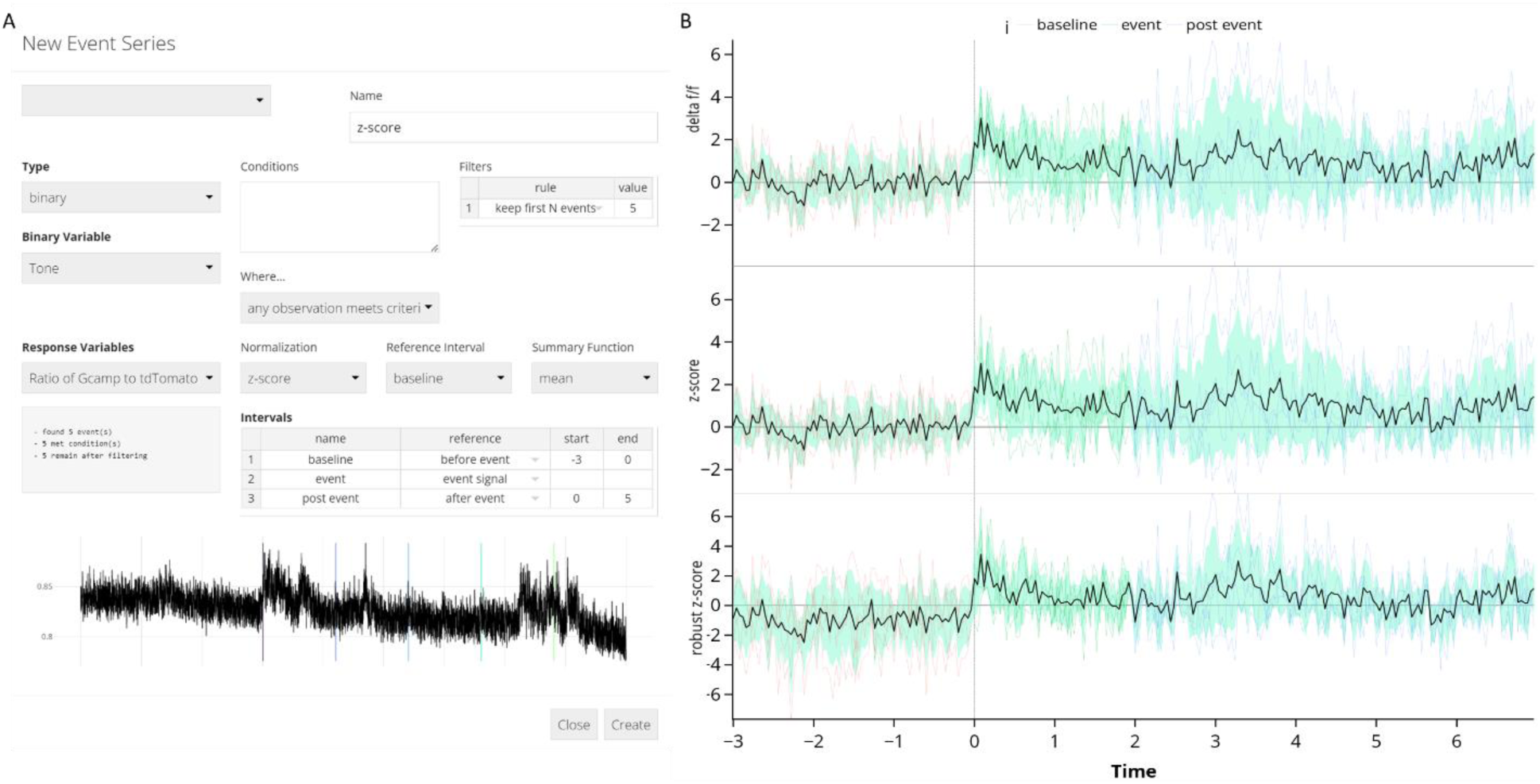
Event-triggered averages for 5 separate tone presentations. (A) Screenshot of events window showing events being created with a 3 second baseline and 5 seconds after the end of the tone. (B) Visualizations of the signal can be made using different normalization methods such as Δ delta F/F, z-score, and the robust z-score. Equations for computing these quantities are in Sec. 2.2.2.

**Figure 5:**
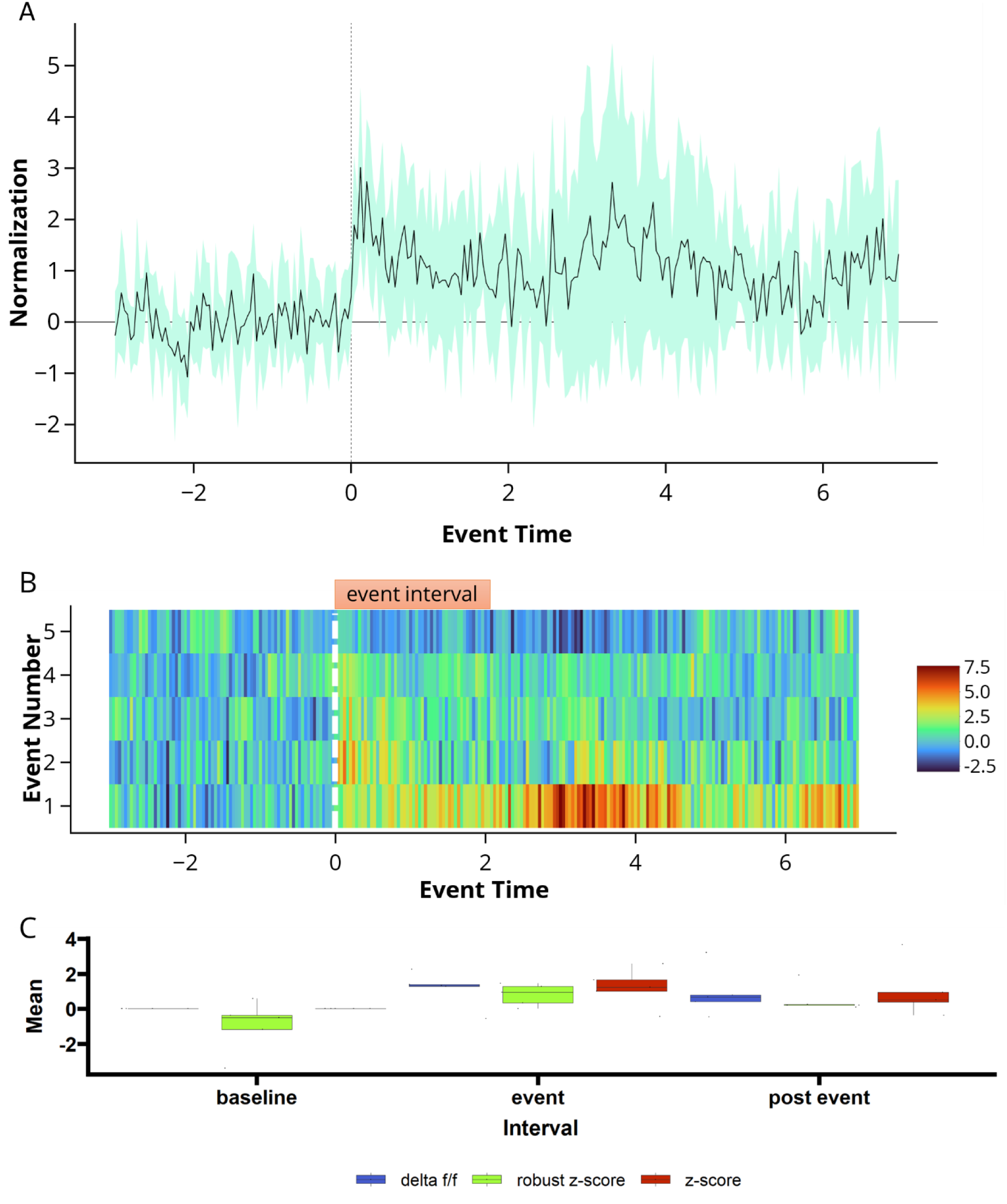
Data analysis options for viewing event-triggered averages. (A) normalized z-score of the data traces of all event-triggered averages calculated during 5 tone presentations. (B) event heatmap of same 5 tone presentation. (C) Interval summaries showing box and whisker plots of the mean values of all events for each interval and normalization scheme.

**Figure 6:**
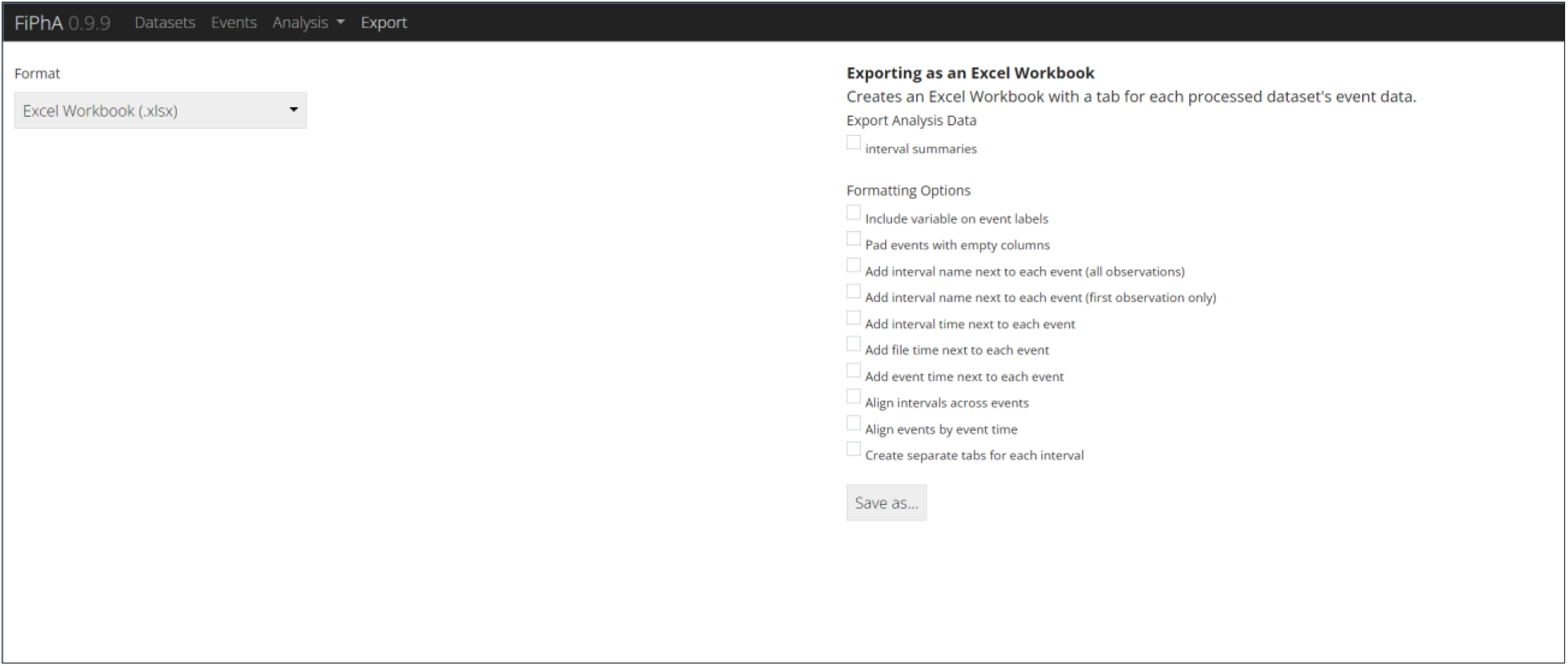
Various export options within the FiPhA app. The user can export as an excel workbook or an R Data Format file to be used for further analysis. The export options enable users to easily copy and paste the data into other graphing software.

**Figure 7:**
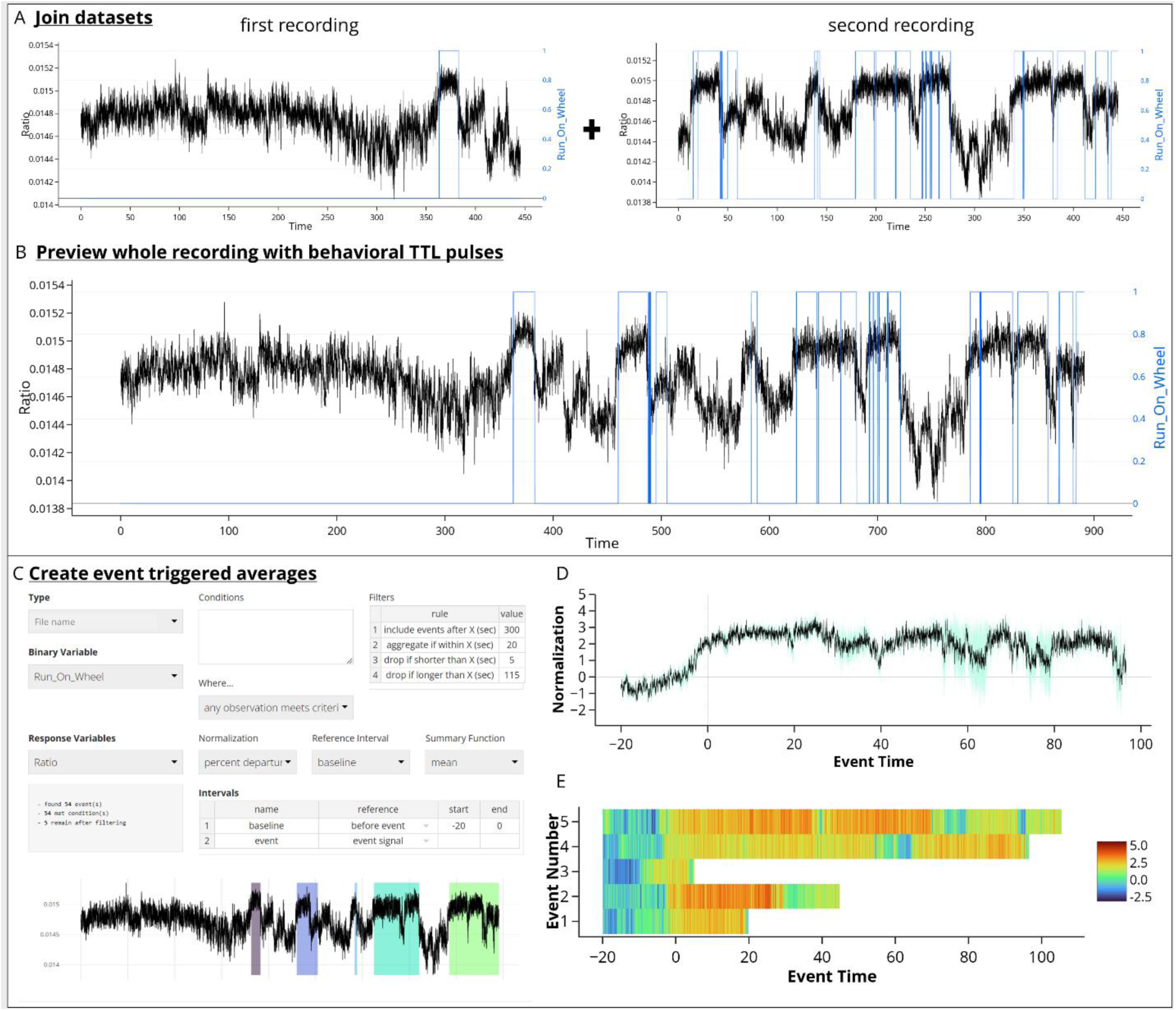
A possible workflow for computing event-triggered averages in the FiPhA app using data collected from a mouse running on a wheel before and after having a drug manipulation. (A) Behavioral tracking (blue) and photometry signal (black) were collected from two separate recordings (before and after injection) and later joined within the FiPhA app (B). (C) Events were then computed with a 20 second baseline and various filtering options, including aggregating smaller running events. Two separate graphs were then created in FiPhA by plotting (D) a normalized and averaged trace graph of all wheel-running events and (E) a heatmap of each normalized running wheel event.

## 4 Discussion

Overall, FiPhA is a flexible and multi-purpose fiber photometry analysis tool that is broadly useful to the field. It is compatible with a broad range of photometry system data types, provides powerful visualization tools and enables customizable event definition and filtering. It is fully open source and developed to meet the needs of the community. While FiPhA was initially developed to provide an integrated approach for spectrally resolved fiber photometry, we have put extensive effort in to making it broadly applicable to various photometry methodologies and data formats. Although, not demonstrated in this manuscript, it will also be useful as the field moves increasingly toward multi-site recordings as processing of multiple photometry data streams simultaneously was built-in early in the design.

Future directions include plans for a standalone R package that allows for integration of FiPhA functionality into user scripts which do not require the interactivity of the Shiny application, as well as the release of FiPhA in an executable format that will be far more user-friendly to those unfamiliar with R and RStudio. We plan to extend features as requested by the community and provide tutorials and training to help increase uptake and usage by interested researchers. Due to its flexibility, wide variety of features, and our plans for continued development and support, we hope that that FiPhA will prove to be of broad utility to the photometry community.

### Animals

All procedures related to animal use were approved by the Animal Care and Use Committee (ACUC) of the National Institute of Environmental Health Sciences. Animal procedures were performed in accordance with the recommendations in the Guide for the Care and Use of Laboratory animals of the NIH.

## Code, Data, and Materials Availability

The code for the package and data for the figures presented in this article is publicly available at https://github.com/mfbridge/FiPhA.

## Acknowledgments

This work was supported by the National Institute of Environmental Health Sciences (ZIC ES103330 to Jesse Cushman) and Social & Scientific Systems, Inc., a DLH Holdings Corp. Company was supported under contract GS-00F-173CA-75N96021F00109.

**Matthew Fletcher Bridge** is a Statistician at Social & Scientific Systems, Inc., a DLH Holdings Corp. Company. He received his BS in Statistics from North Carolina State University in 2014, and has an academic background in chemistry, physics, and computer science. His research interests include simulation and algorithm development, mathematical modeling of biologic processes, and data visualization. He is a member of the American Statistical Association and Society for Neuroscience.

**Leslie Rae Wilson** is a Research Scientist in the Neurobehavioral Core at NIEHS. She received her PhD in Chemistry from North Carolina State University in 2018. Her current research interests include measuring neurochemicals in the brain using various techniques and linking those signals with specific behavioral outcomes. She is a member of the Society for Neuroscience (SfN), the Pavlovian Society, and the Organization for the Study of Sex Differences.

**Sambit Panda** received a BS from the Department of Biomedical Engineering (BME) at the Joint Department of Biomedical Engineering at University of North Carolina and North Carolina State University in 2018. He then went to Johns Hopkins University where he received a M.S.E. degree from the Department of Biomedical Engineering at Johns Hopkins University (JHU) in 2020. He is currently a Ph.D. candidate in the Department of Biomedical Engineering at JHU. He is also affiliated with the Institute of Computational Medicine. He does biomedical data science, primarily focusing on high-dimensional hypothesis testing, causal inference, and behavioral neuroscience.

**Korey Daniel Stevanovic** is a Research Scientist in the Neurobehavioral Core at NIEHS. He received his M.Sc. in Neuroscience and Education from Columbia University in 2009. His current interests include methods development and the topics of motivation and addiction. He is a member of the Society for Neuroscience (SfN), the Pavlovian Society, and the Organization for the Study of Sex Differences.

**Ayland Cash Letsinger** is a Postdoctoral Fellow in the Neurobehavioral Laboratory at the NIEHS. He received his PhD in Exercise Physiology from Texas A&M University in 2019. His research aims to discover the neural foundations that determine habitual wheel-running behavior, with a specific emphasis on characterizing neurotransmitter and modulator activities using fiber photometry. He is an active member of the American College of Sports Medicine.

**Sandra McBride** is a Senior Statistician at Social & Scientific Systems, Inc., a DLH Holdings Corp. Company. She received her PhD in Statistics from Stanford University in 2000 and did post doctoral training in Bayesian statistics at Duke University. Her research interests span spatial and time series modeling, point process models, survival models and mixed models.

**Jesse Daniel Cushman** received his PhD in the lab of Dr. Michael Fanselow in 2010 and did post doctoral training in the lab of Dr. Mayank Mehta. He ran the UCLA Behavioral Testing Core for three years before starting the Neurobehavioral Core Laboratory at the National Institute of Environmental Health Science in 2016. His primary interests are in behavioral and neurophysiological analysis of hippocampal function.

